# Response of siliceous marine organisms to the Permian-Triassic climate crisis based on new findings from central Spitsbergen, Svalbard

**DOI:** 10.1101/2023.09.01.555975

**Authors:** W.J. Foster, G. Asatryan, S. Rauzi, J. Botting, S. Buchwald, D. Lazarus, T. Isson, J. Renaudie, W. Kiessling

## Abstract

Siliceous marine ecosystems play a critical role on the Earth’s climate system through its influence on organic carbon burial and rates of marine authigenic clay formation (i.e. reverse weathering). The ecological demise of silicifying organisms associated with the Permian-Triassic mass extinction is postulated to have elevated rates of marine authigenic clay formation, resulting in a prolonged greenhouse climate during the Early Triassic. Yet, our understanding of the response of siliceous marine organisms during this critical interval is poor. Whilst radiolarians experienced the strongest diversity loss in their evolutionary history and perhaps also the greatest population decline of silica-secreting organisms during this event, only a small number of Griesbachian (post-extinction) localities that record siliceous organisms are known. Here, we report newly discovered latest Changhsingian to early Griesbachian (*Clarkina meishanensis* - *Hindeodus parvus* Zone) radiolarians and siliceous sponge spicules from Svalbard. This fauna documents the survival of a low-diversity radiolarian assemblage alongside stem-group hexactinellid sponges making this the first described account of post-extinction silica-secreting organisms from the Permian/Triassic boundary in a shallow marine shelf environment and a mid-northern palaeolatitudinal setting. Our findings indicate that latitudinal diversity gradients for silica-secreting organisms following the mass extinction were significantly altered, and that silica productivity was restricted to high latitude and deep water thermal refugia. This result has potential to further shape our understanding of changes to marine porewater and seawater dissolved silica levels and in turn rates of reverse weathering, with implications for our understanding of carbon cycle dynamics during this interval. This also suggests that the export of organic carbon to the deep ocean was not as severely impacted at non-equatorial latitudes.

**Key Points:**

- We document the first occurrence of siliceous sponge spicules and radiolarians (biogenic silica) from a mid-northern paleolatitude following the mass extinction event
- Holdover radiolarian species show poleward range shifts
- The ecological composition and the restriction to shallow water oxygenated facies suggests a shallow mid-latitude refuge for siliceous marine organisms
- This result has potential to further shape our understanding of changes to marine dissolved silica levels and in turn rates of reverse weathering, with implications for our understanding of Permian-Triassic carbon cycle dynamics.

## 1 Introduction

The Permian-Triassic mass extinction was the most catastrophic extinction event of the Phanerozoic, resulting in major changes in marine ecosystem diversity, ecosystem functioning and the biosphere’s evolutionary pathway (Muscente et al., 2018). The mass extinction was notably catastrophic for biosiliceous productivity, resulting in a biodiversity crisis for silica-secreting organisms (Liu et al., 2013; De Wever et al., 2006) and the Early Triassic chert gap, a five-million-year interval of global low chert occurrence in the global rock record (Isozaki, 1997; Racki, 1999; Beauchamp and Grasby, 2012; Isson et al., 2022; Yang et al., 2022). Notably, the Early Triassic chert gap directly deviates from observations of increased chert burial following other Phanerozoic hyperthermals (e.g., PETM; Triassic-Jurassic) as a result of increased silicate weathering rates (Kump, 2018; Ritterbush et al., 2014; Ritterbush 2019; Isson et al., 2020).

It is traditionally viewed that following a carbon-injection event, an increase in atmospheric *p*CO_2_ would foster an increase in silicate weathering and in turn carbon burial through the deposition of carbonate and chert, which would act to reduce *p*CO_2_ levels to pre-injection conditions on the ∼10^5^ year timescale (Urey, 1952, Walker et al., 1981, Berner et al., 1983; Isson et al., 2020). The Permian-Triassic hyperthermal deviates from this framework of climate regulation. Here, climate recovery was slow, with temperatures remaining elevated for millions of years in the aftermath of the mass extinction. This has led to the view of a fundamentally altered climate system in the wake of the Permian-Triassic mass extinction, in which silicate weathering was ineffective at drawing down *p*CO_2_ and lowering temperatures (Kump, 2018; Isson et al., 2022). It is postulated that volcanic carbon release may have been substantially large enough to ‘exhaust’ the supply of silicate minerals available for weathering at Earth’s surface (Kump, 2018). Recent work has also called upon the widespread loss of marine silicifying organisms, and resulting increase in marine dissolved silica concentration, as a catalyst for increased CO_2_ release via marine authigenic clay formation (Isson et al., 2022). Marine authigenic clay formation consumes alkalinity and dissolved cations initially released via the process of silicate weathering, and in turn acts to re-liberate the CO_2_ initially captured as dissolved carbon (HCO_3-_). Overall, marine authigenic clay formation acts to recycle carbon within the ocean-atmosphere system, variation in the extent of this process can drive changes in atmospheric *p*CO_2_ levels and global climate (Isson & Planavsky, 2018). In this view, the demise of silicifying ecosystems (and the Early Triassic chert gap) are directly responsible for the anomalous Triassic warm period (Isson et al., 2022).

Estimates of changes in marine dissolved silica concentration used in global carbon-silica cycle models are reliant on direct observations of biogenic silica burial in the global rock record (Isson, et al., 2022). Therefore, finding new Permian/Triassic successions that record siliceous organisms and documenting the extent of biogenic silica productivity in different environments is essential in understanding the evolution of silica-secreting organisms, their refugia during climate crises, and impact on changes to global climate.

Siliceous sponges and radiolarians, both of which evolved in the latest Precambrian, were the predominant silicifying organisms during the Permian-Triassic. Globally, radiolarians, along with acritarchs, were the primary fossil record of plankton biodiversity through the Permian-Triassic transition and suffered from the most severe extinction in their evolutionary history (De Wever et al., 2006; O’Dogherty et al., 2010). Many aspects of this extinction, and the subsequent radiation are poorly known due to the paucity of material in the earliest Triassic (O’Dogherty et al., 2010). Furthermore, post-extinction (Griesbachian) radiolarians have only been identified in two regions in the world, New Zealand (Takemura et al., 2002; Hori et al., 2011) and Japan (Sano et al., 2010) (Fig. 1). Siliceous sponges show a similar trend, where they thrived during the Permian, depositing large quantities of biogenic chert across large areas of shelf and slope settings (Beauchamp and Baud, 2002), and then virtually vanish following the extinction event. Only a single morphology and rare occurrence of undescribed smooth siliceous monaxone sponge spicules have been reported from the Griesbachian in South China (Liu et al., 2013) and Canada (Grasby and Beauchamp, 2009) (Fig. 1), but in both cases it is possible that these post-extinction sponge spicules were reworked from pre-extinction sediments (Beauchamp, pers. comm). This lack of known localities with silica skeletal remains from the Griesbachian, and even the subsequent Dienerian and Smithian, suggests that the absence of biogenic chert deposits carries a genuine biological (productivity) signal.

**Figure 1.**
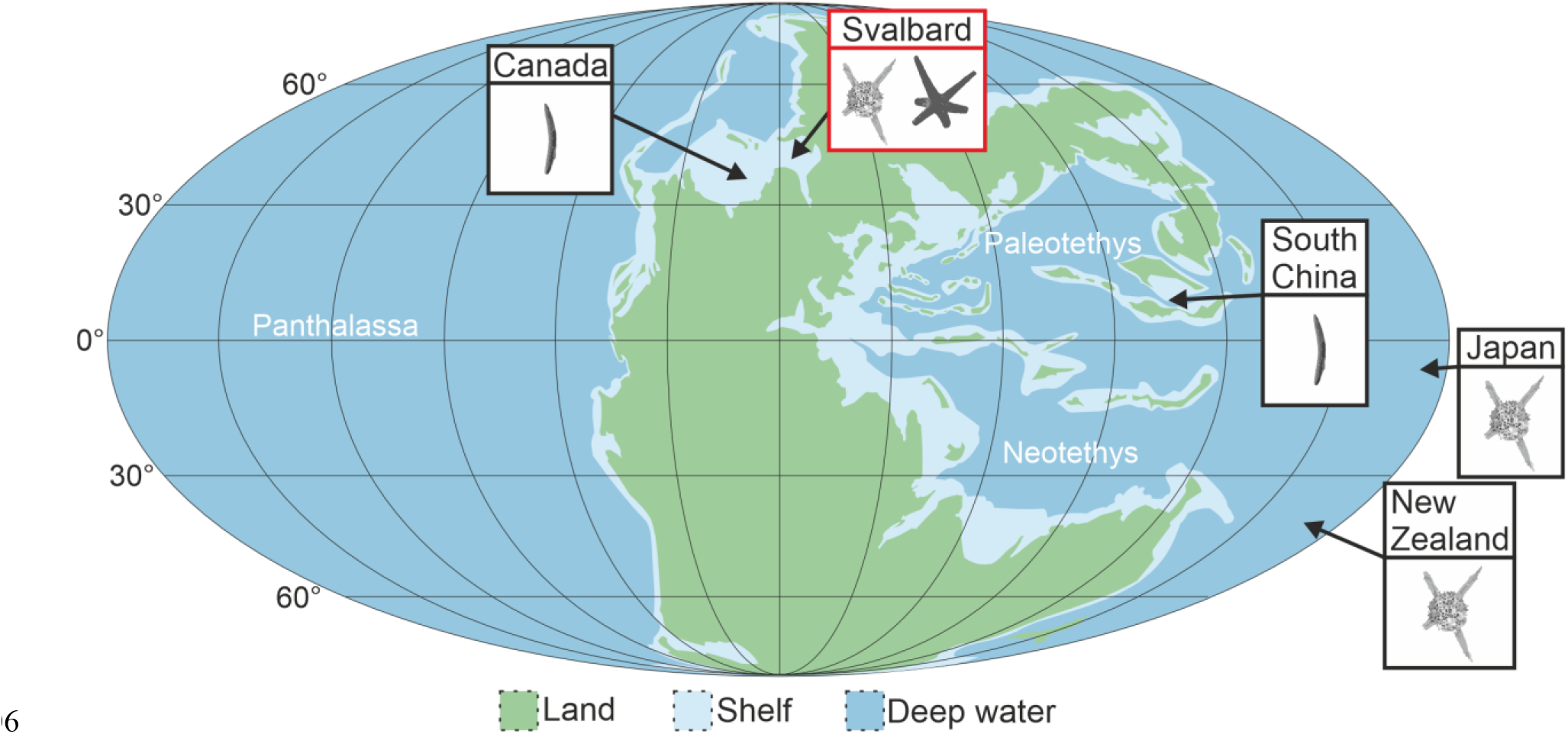
Paleogeographic map for the Permian/Triassic boundary showing the records of radiolarians and siliceous sponges in the Griesbachian, Early Triassic. Svalbard (45°N), this study; Canada (40°N) (Grasby and Beauchamp, 2009); South China (8°N), (Liu et al. 2013); Japan (6°S) (Sano et al. 2012); and New Zealand (36°S), (Takemura et al. 2002). Base map after Blakey (2012).

Here we report new finds of radiolarians and sponge spicules in the Vikinghøgda Formation at Lusitaniadalen and Deltadalen, Svalbard (Fig. 2). These findings represent the first described post-extinction (Changhsingian-Griesbachian) record of siliceous sponges and radiolarians from both the Boreal Realm and a shallow depositional setting, with implications for improving our understanding of the distribution of post-extinction silica-secreting organisms, and refugia during global warming events.

**Figure 2.**
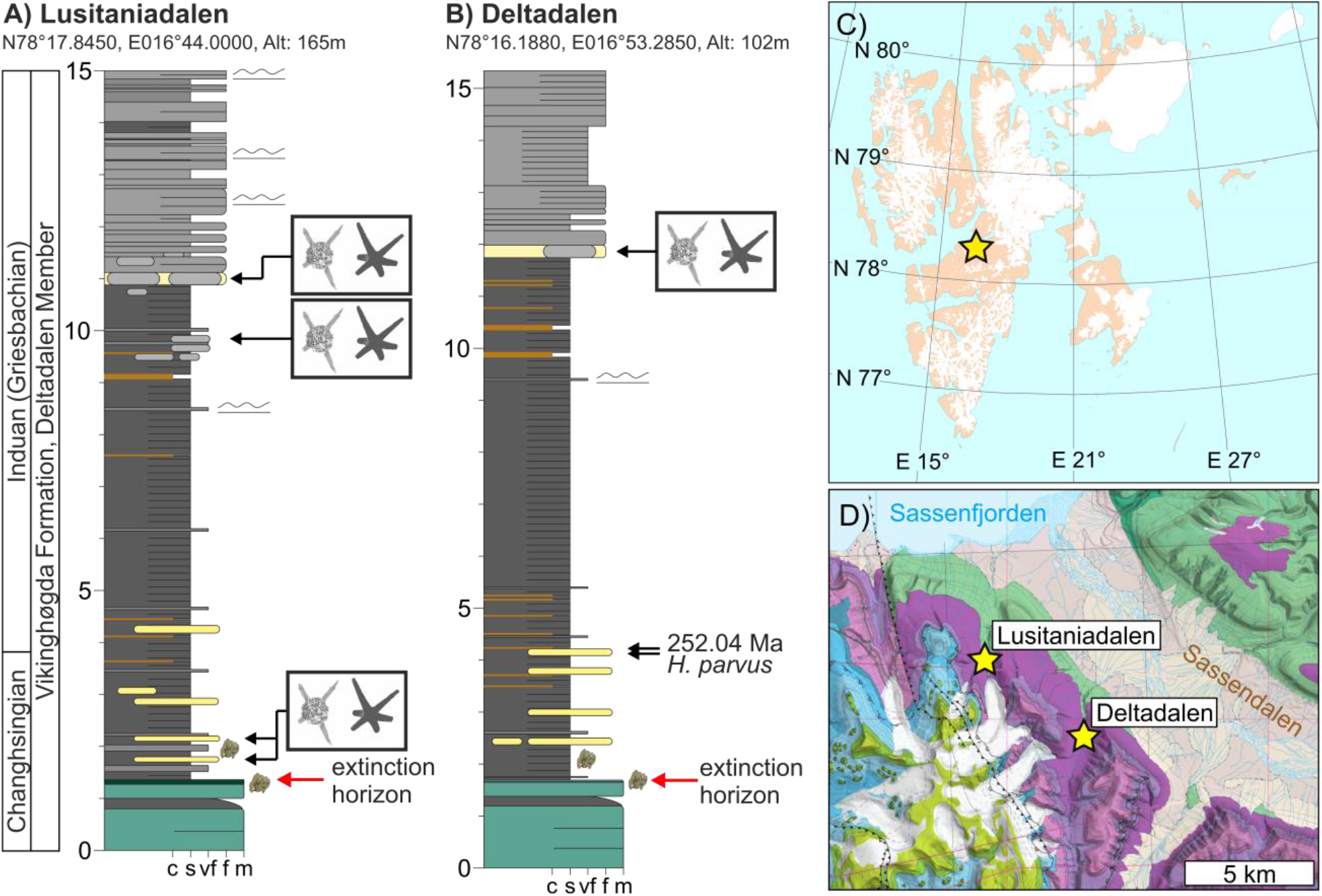
Stratigraphic succession at Lusitaniadalen and Deltadalen, Svalbard. A) Radiolarians and sponges recorded at 1.8, 9.8 and 11.0 m into the Vikinghøgda Formation at Lusitaniadalen (78.2986, 16.7331) and B) 12.0 m into the Vikinghøgda Formation at Deltadalen (78.2698, 16.8881), Svalbard. The samples that yielded siliceous fossils are shown by the avatars. Position of the Permian/Triassic boundary and the U/Pb date comes from Zuchuat et al. (2020). C) Inset map of Svalbard. D) Geological map of Sassendalen. Geological map after https://geokart.npolar.no/geologi/GeoSvalbard.

## 2 Materials and Methods

### 2.1 Geological Setting

The Vikinghøgda Formation can be traced across central and eastern Svalbard and records deposition through the latest Permian and entire Early Triassic in a mid-paleolatitude (∼45°N, Hounslow et al., 2008), siliciclastic, open-marine, shelf setting (Mørk et al., 1999). Both Lusitaniadalen (78.2986, 16.7331) and Deltadalen (78.2698, 16.8881), record the Permian-Triassic transition and have been stratigraphically-constrained through magnetostratigraphy, U-Pb geochronology and biostratigraphy. The radiolarians and sponges from these sections are, therefore, well-constrained as belonging to the latest Changhsingian to early Griesbachian *Clarkina meishanensis* -*Hindeodus parvus* Conodont Zones (Mørk et al., 1999; Zuchuat et al., 2020) (Fig. 2). The concretions come from interbeds of siltstones and very fine sandstones, where the heterolithic beds are bioturbated by shallow burrowing *Skolithos* and *Arenicolites* and both record ripple structures and a diverse fauna, which indicate deposition in an oxygenated, shallow, marine setting (Foster et al., 2017; 2022).

### 2.2. Materials and sample processing

Samples were collected every 50 cm throughout the succession from all the lithologies. Each sample was then disaggregated using 10% hydrogen peroxide with the solution changed every 48 h. In addition, samples were collected from each concretionary horizon. The samples from the concretionary horizons were mechanically disaggregated into 1–2 cm-sized blocks, and the pieces that did not have fossils on their surfaces were disaggregated with the buffered formic acid technique (Jeppsson and Anehus, 1995). To maximize yield, the residue was collected at c. 12 hour intervals, washed thoroughly with tap water to remove any excess solution and to avoid crystal growth, and dried. The fossils were then picked using standard microfossil picking techniques and mounted onto stubs for imaging using a scanning electron microscope. The specimens are housed in the University of Oslo, Natural History Museum.

### 2.3. Analyzing species diversity

To assess the differences in species richness between the samples from Svalbard and between the different radiolarian localities for the Griesbachian, a sample-size-based rarefaction using absolute abundance data was done for the samples from Svalbard and a coverage-based rarefaction curve using incidence data was done for all of the Griesbachian radiolarian samples using the data from Sano et al., (2010; 2012), Hori et al., (2011), and Takemura et al., (2002). Estimates of the asymptote species richness for the samples from Svalbard was done using the Chao 1 richness index (Chao et al., 2009). The rarefaction curves and Chao index were computed using the iNEXT package (Hsieh et al., 2014) in R.

## 3 Results

Sponge spicules and radiolarians were discovered in the concretionary horizons of the Vikinghøgda Formation indicated in Figure 2. The fauna identified from the Vikinghøgda Formation are not considered to have been reworked from the underlying pre-extinction, spiculitic Kapp Starostin Formation for the following reasons: (1) the Kapp Starostin Formation and the Vikinghøgda Formation in central Spitsbergen record stark lithological differences and lithological features of the Kapp Starostin Formation are not recorded in association with the siliceous fossils (e.g., glauconitic sand), (2) the fossils are well-preserved with delicate thin skeletal elements that would have been destroyed if reworked, (3) abrasion on some specimens is interpreted as a consequence of the disaggregation methods, (4) the Vikinghødga Formation does not record significant bioturbation that could have reworked the fossils into younger strata, and (5) the silicified fauna is dominated by Mesozoic taxa.

The identified fauna includes 238 radiolarians, of which 87 could be identified to the genus-level and 61 to the species-level. Ten distinct species were identified (Fig. 3) and only four were Permian holdovers (*Entactinia itsukaichisensis, Hegleria mammilla, Hegleria* sp., *Grandetortura nipponica*). Five species are recorded from the Griesbachian for the first time, including one new species (*Entactinia* n.sp.). Three of those were previously known from the Early Triassic but represent range extensions back to the *H. parvus* Zone. In addition to no signs of reworking, these Triassic species support that the radiolarian assemblage does not represent a reworked Permian fauna. One species is recorded in the Triassic for the first time (*Grandetortura nipponica*), which records one of the last occurrences of the order Latentifistularia that subsequently went extinct in the Pelsonian (Feng et al., 2000).

**Figure 3.**
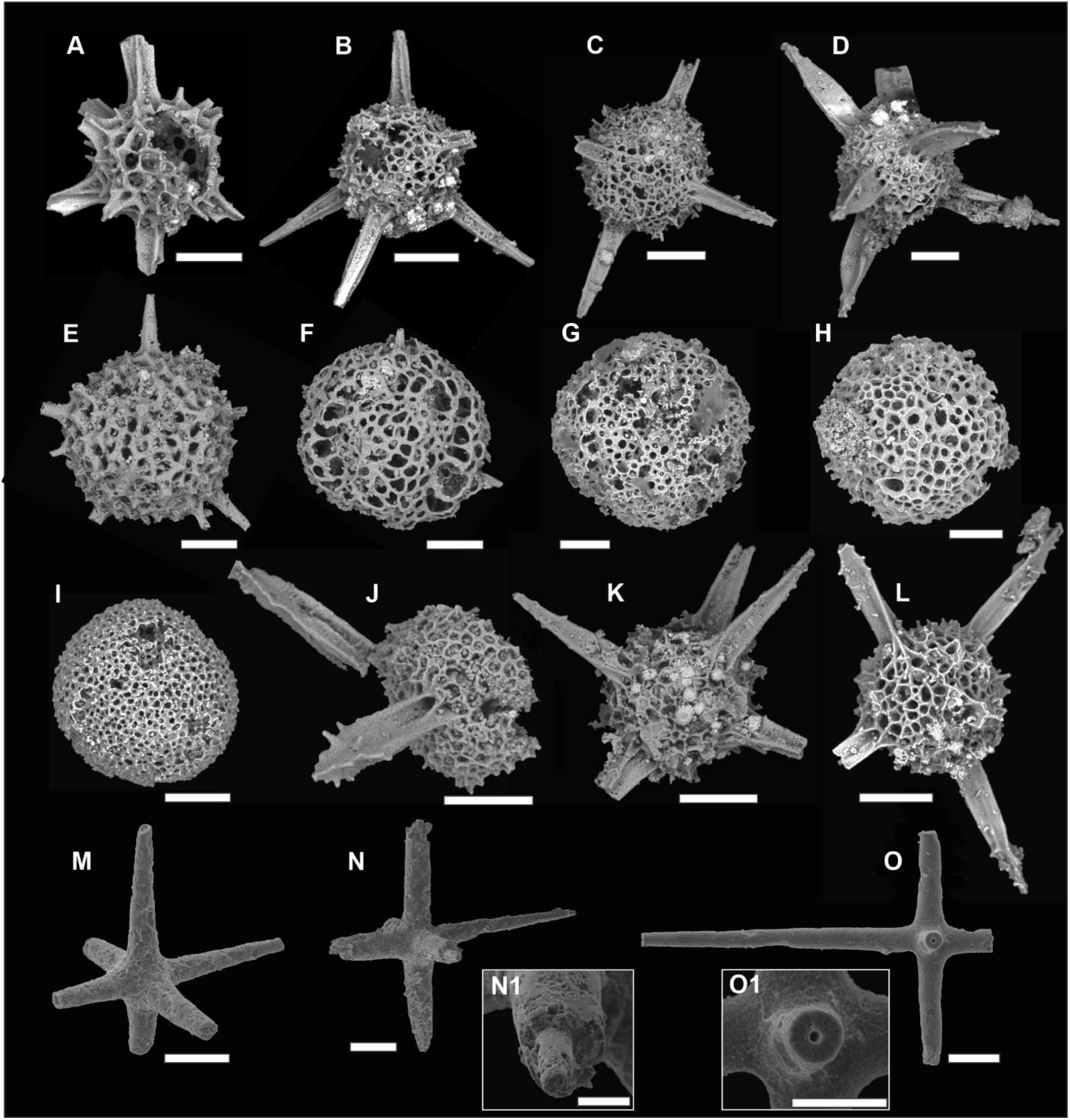
Radiolarians and sponge spicules from the Griesbachian (*Hindeodus parvus* Conodont Zone) of central Svalbard. A) *Entactinia itsukaichisensis* (Sashida and Tonishi 1985), 1JB1.1. B) *Entactinia nikorni* (Sashida and Igo 1992), 1JB2.1. C) *Entactinia* cf. *chiakensis* (Sashida and Igo 1992), JB4.19. D) *Entactinia* n. sp., 1JB2.4. E) *Polyentactinia* cf. *phattalungensis* Sashida and Igo 1992, JB5.19. F) *Grandetortura nipponica* Sashida and Tonishi 1991, 2JB8.4. G) *Hegleria mammilla* (Sheng and Wang 1985), 1Rad2.6. H) *Hegleria* sp., 1RAD3.7. I) Gen et sp. indet., 1Rad4.3. J) *Thaisphaera*? *igoi* Kamata 1999, 2JB9.10. K) *Thaisphaera*? *igoi* Kamata 1999, 2JA5.5. L) *Thaisphaera*? *igoi* Kamata 1999, 1Rad3.6. M-O) Hexactinellida sponge spicules, N1 and O1 close-up views of the axial canal. scale bar = 100 μm, except N1 and O1 where scale bar = 30μm.

The observed species richness of the samples from Svalbard is highly dependent on sample size and a sample-based rarefaction curve for the three samples from Svalbard (Fig. 4) suggest that the asymptote for species richness from these assemblages has not yet been reached and that the samples are likely more species-rich than observed. Comparisons of species richness between Svalbard and other Griesbachian radiolarian localities could only be made using a coverage-based rarefaction which suggest that the asymptote for species richness has not yet been reached for any Griesbachian locality (Fig. S2B) and the samples from Svalbard are not significantly different from the species richness of the other localities.

**Figure 4.**
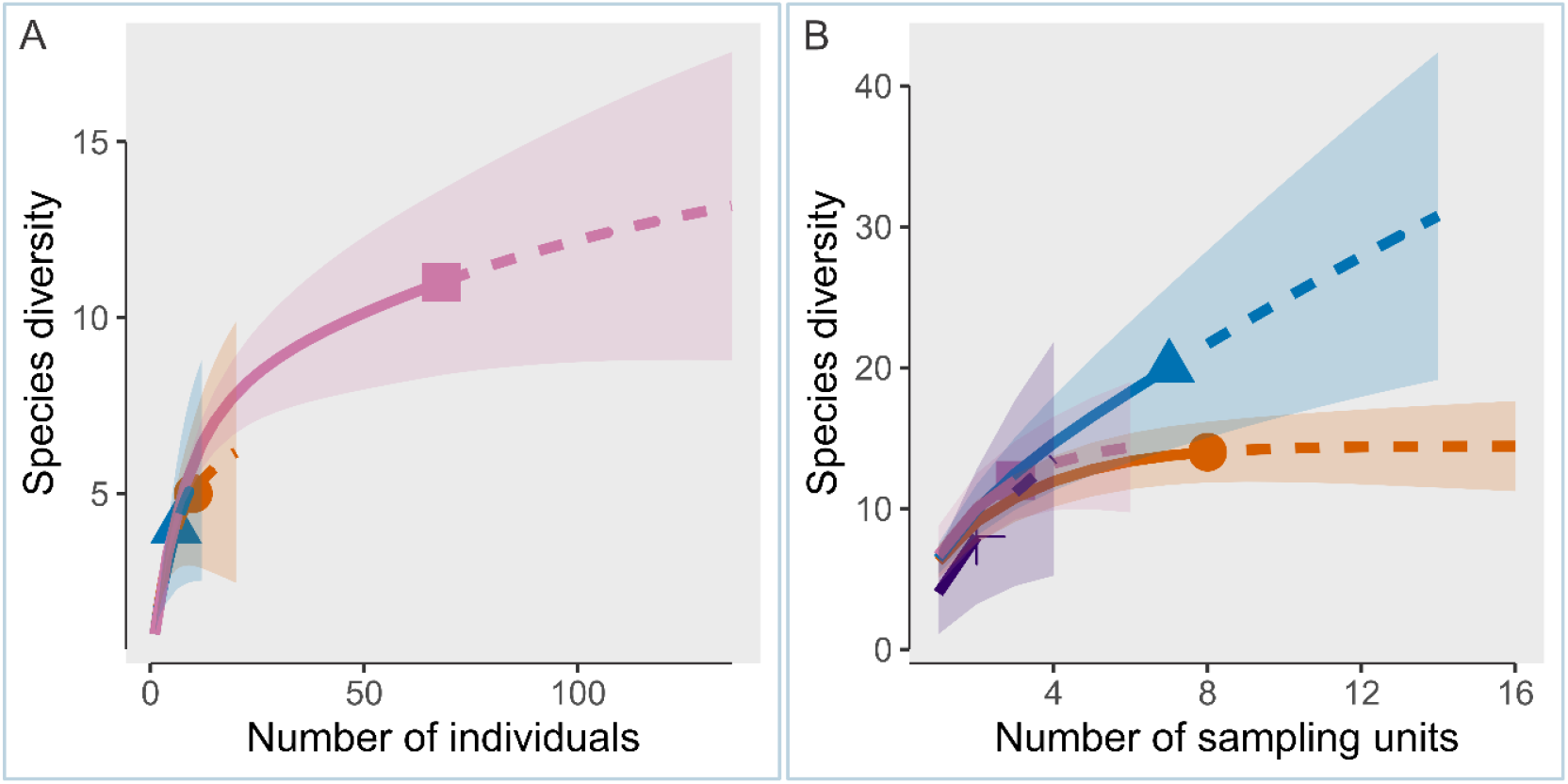
Rarefied species richness curves (solid line segment) and extrapolation (dotted line segments) sampling curves with 95% confidence intervals (shaded areas) for the Griesbachian Radiolarians. A) Sample-size-based rarefaction for the samples from Svalbard (this study), from 11.8 m into the Vikinghøgda Formation at Deltadalen (pink square), 9.8 m (orange circle), and 11.0 m (blue triangle) at Lusitaniadalen. B) Coverage-based rarefaction curve for the Griesbachian radiolarians collected from Arrow Rocks (orange circle) and Waiheke Island, New Zealand (purple cross); Mino Terrane, Japan (blue triangle); and the Vikinghøgda Formation in Svalbard (pink square). Data: Svalbard = This Study, Mino terrane, Japan = Sano et al. (2010; 2012), Waiheke Island, New Zealand = Hori et al., (2011), Arrow Rocks, New Zealand = Takemura et al., (2002).

The fauna also includes abundant siliceous sponge spicules that can be confidently described as oxyhexactin, which are siliceous spicules with six unbranched rays perpendicular to one another that taper to a point (Boury-Esnault and Rutzler, 1997). The lack of any other siliceous spicule morphology suggests both a low diversity and that the spicules belong to a stem-hexactinellid, representing a Permian holdover. In addition, a thin section from the first concretionary horizon above the mass extinction event at Lusitaniadalen, also records radiolarians and oxyhexactin spicules in four thin sub-beds (up to 2 cm).

The radiolarian and sponge fossils are also interpreted to represent para-autochthonous assemblages. This is because the fossils are reported from the base of the Vikinghøgda Formation which is characterized by silty mudstone and deposition in a low-energy environment (Mørk et al., 1999). In addition, the composition of the radiolarians represent a typical shallow water assemblage (see Discussion). The fossils are, therefore, likely to have undergone some spatial-averaging but their exquisite preservation also suggests that they were rapidly buried. Notably, the silicified fossils from the base of the Vikinghøgda Formation with 3D preservation are restricted to concretionary beds (see also Foster et al., 2017), whilst macrofossils in the surrounding silts and muds are typically flattened moulds. The radiolarians and sponge spicules found here also record exquisite 3D preservation and have only been observed within concretionary beds. This suggests that the fossils were preserved within the concretions during early diagenetic stage that prevented the siliceous fossils from undergoing diagenetic dissolution or partial-dissolution, as observed in the underlying Kapp Starostin Formation.

## 4 Discussion

Numerous post-extinction refugia have been suggested from various paleogeographic locations and depositional settings (Beatty et al., 2008; Twitchett et al., 2004; Godbold et al., 2017) and in particular shallow marine settings in the Boreal realm (Wignall et al., 1998; Beatty et al., 2008). Our new data highlight that the fossil record from Svalbard is exceptional in not only recording diverse and complex marine ecosystems (Foster et al., 2017), but also recording faunal groups currently unknown from coeval shallow marine settings, including radiolarians (this study), hexactinellid sponges (this study), large bivalves (Foster & Buchwald, pers. obvs.), red algae (Wignall et al., 1998) and bryozoans (Nakrem and Mørk, 1991). In addition, the only other reports of Griesbachian siliceous sponge spicules come from South China and Canada (Grasby and Beauchamp, 2009, Liu et al., 2013). Sponge spicules are reported as occurring in Wadi Maqam (Oman) during the Griesbachian in a review by Isson et al. (2022), but this occurrence could not be verified. The absence of these groups recorded in Svalbard from intensely studied shallow marine successions from the Tethys and Panthalassa oceans suggests that their absence in these equatorial settings represents a biological signal and more habitable conditions for siliceous and calcareous organisms in the Boreal Realm during the Permian-Triassic climate crisis.

One explanation for the absence of siliceous marine organisms from low-latitude, shallow marine localities could be thermal stress associated with the mass extinction. Species track their thermal niches through range shifts resulting in latitudinal diversity gradients in marine species (including primary planktic production) becoming significantly altered; with an equatorial depression in diversity and increased diversity towards temperate latitudes (Yasuhara et al., 2020; Chaudhary et al., 2021). Consistent with this observation, the altered latitudinal diversity gradient during the Griesbachian implies a relatively greater ecological impact in the tropics (Foster and Twitchett, 2014; Song et al., 2020). However, mid-to high-latitude regions are hypothesized to have higher turnovers in faunal composition, owing to the combination of local extinctions, extirpations and poleward range extensions of low-latitude taxa (Reddin et al., 2022). The range extensions and dominance of many Triassic fossil species and genera to the Griesbachian belonging to new, post-extinction orders because of new data from Svalbard (this study, see also Foster et al. 2017) also suggests high extinction and origination rates in this region.

The Permian holdover radiolarians *Hegleria mammilla, Entactinia itsukaichisensis*, and *Grandetortura nipponica* recorded in this study all show poleward latitudinal range extensions out of the tropics across the mass extinction event. In addition, laboratory cultures show that subtropical radiolarians can only survive for a few days when exposed to high temperature stress (Andersson et al., 1989), and the overall low-diversity recorded in Svalbard could be attributed to thermal stress even at higher latitudes. This is in line with predictions of climate warming, oxygen loss and metabolic theory, and the observation that the toll of the extinction event for marine genera was greater in higher latitudes than in the tropics (Penn et al., 2018). Taken together, the impact of climate change would have been detrimental to higher latitude ecophysiotypes, yet the mid-to high-latitude Boreal Ocean setting of Svalbard must still have had substantially lower sea-surface temperatures and higher rates of primary productivity compared to the tropics. Moreover, a strong increase in primary productivity is recorded in the latest Changhsingian and early Griesbachian at Lusitaniadalen by enhanced input of chlorophyll-derived biomarkers (Buchwald, pers. obvs.). With radiolarian abundance being linked to chlorophyll concentrations (Lampitt et al., 2009), food availability was likely increased for the radiolarian assemblage, which possibly contributed to the relatively high radiolarian abundance in this setting.

An alternative adaptation to thermal stress is to track a thermal niche through migrating down the water column to cooler waters. Our radiolarian assemblage is dominated by entactinarians, two latentifistularians and a single spumellarian, which, consistent with the facies interpretation (Foster et al., 2017), is indicative of a shallow water assemblage (Xiao et al., 2017). This also supports the interpretation that the radiolarians represent a para-autochthonous assemblage. For the silica-secreting organisms in Svalbard, however, the water column migration of taxa would not have been possible as the deeper water facies are inferred to have been anoxic, or euxinic, for most parts of the Griesbachian (Nabbefeld et al., 2010; Zuchuat et al., 2020). Siliceous organisms in Svalbard were, therefore, restricted to shallow marine settings and only groups that could tolerate the associated environmental stressors in this setting would survive. The restriction of diverse marine fossil assemblages to shallow marine ecosystems was also demonstrated with the distribution of trace fossils in NW Pangaea (Beatty et al., 2008). The virtual absence of the Albaillellaria and Latentifistularia from the Svalbard samples also suggests that these orders were unable to respond to expanding oxygen minimum zones by migrating into shallower settings. It is therefore clear that consistent with recent studies (e.g., Penn et al., 2018) the combined effects widespread anoxia and high sea surface temperatures played a key role in limiting the diversity and abundance of silica-secreting organisms and primary production.

The low species richness of siliceous organisms in Svalbard could be related to the shallow depositional setting rather than a consequence of the climate crisis. Modern hexactinellids for example generally inhabit deep-water habitats (greatest diversity at 300-600 m depths) and only a few populations inhabit shallow settings (Leys et al., 2007). The distribution of sponges during Permian was, however, more widespread with sponges being abundant in very shallow, inner ramp settings, which also lead to the development of glass ramps along the NW Pangaea coastline (Gates et al., 2004) and extensive chert deposition in shallow basins on the South China block (Liu et al., 2013). In addition, radiolarians are recorded as occupying water depths from 100s to 1000s of meters deep with a distinct ecological zonation in the Permian (Xiao et al., 2017). This may also explain why the oceanic settings of Japan and New Zealand both record a greater diversity of radiolarians in the Griesbachian, which includes latentifistularians, albaillellarians, spumellarians, entactinarians and nassellarians (see Table S1), i.e. these assemblages will have much larger spatial averaging of habitats. Alternatively, the shallow epicontinental setting of the Barents Sea -Svalbard area compared to the open oceanic settings of Panthalassa may explain the paucity of both pre- and post-extinction radiolarians assemblages. The low species richness of radiolarians and hexactinellids from Svalbard does not, therefore, negate the mid-latitudinal setting as a refuge for shallow water forms.

The preferential survival of siliceous organisms in mid-latitude and deep-water habitats supports the view that thermal stress drove the collapse of silica productivity during the climate crisis and provides a further constraint on the extent of biogenic silica burial decline following the end-Permian mass extinction. This restriction of biogenic silica productivity has key implications for the regulation of Earth’s climate during the Griesbachian. Directly, radiolarians are major exporters of organic carbon to the deep ocean (Lampitt and Johns, 2009), and the widespread decline in large-scale silica-production would have limited the oceans ability to sequester carbon dioxide from the atmosphere. Further, the widespread loss of silicifying organisms potentially increased marine dissolved silica concentration and marine authigenic clay formation, trapping the Earth in a prolonged hyperthermal state (Isson et al., 2022). The findings of siliceous organisms and thin siliceous beds in Svalbard (this study) and potentially also Canada (Grasby and Beauchamp, 2009) suggests a poleward expansion of silica productivity.

Observations of radiolarians and sponge spicules here suggest a poleward expansion of silica productivity and spatially varied changes in biogenic silica burial following Permian-Triassic extinction event. This has potentially critical implications for our understanding of both global marine dissolved silica levels and also local porewater conditions (the locus of marine clay formation). For instance, biogenic silica fixation and in turn local scale depletion of seawater dissolved silica could have been most intense at mid-latitude and deep-sea environments (distinct to the modern day and Permian systems). Relatively elevated biogenic silica deposition in these restricted environments may further alter porewater dissolved silica levels. Overall, this shift in the locus of biogenic silica production and deposition has potential to influence the global distribution and rates of marine clay formation and in turn atmospheric *p*CO_2_ levels.

## Acknowledgments

We would like to thank Hans Arne Nakrem for his support in the search for invertebrate communities from the Permian-Triassic transition of Svalbard. We also thank Benoit Beauchamp and Michael Hautmann for their comments on an earlier version of this manuscript. WJF is funded by the Deutsche Forschungsgemeinschaft (Project No. FO1297/1-1). This work is a contribution to the Research Unit TERSANE (FOR 2332: Temperature-related stressors as a unifying principle in ancient extinctions). We acknowledge financial support from the Open Access Publication Fund of Universität Hamburg.

## Open Research

All of the data used in this study is available in the supplementary files and the specimens have been accessioned at the natural History Museum, University of Oslo.

## References

Anderson, O.R., Bennett, P., & Bryan, M. (1989), Experimental and observational studies of radiolarian physiological ecology: 3. Effects of temperature, salinity and light intensity on the growth and survival of Spongaster tetras maintained in laboratory culture. Marine Micropaleontology, 14, 275–282. 10.1016/S0377-8398(96)00011-4

Beatty, T.W., Zonneveld, J.P., & Henderson, C.M. (2008), Anomalously diverse Early Triassic ichnofossil assemblages in northwest Pangea: a case for a shallow-marine habitable zone: Geology, 36, 771–774. 10.1130/G24952A.1

Beauchamp, B., & Baud, A. (2002), Growth and demise of Permian biogenic chert along northwest Pangea: evidence for end-Permian collapse of thermohaline circulation: Palaeogeography, Palaeoclimatology, Palaeoecology, 184, 37–63. 10.1016/S0031-0182(02)00245-6

Berner, R. A., Lasaga, A. C. & Garrels, R. M. Carbonate-silicate geochemical cycle and its effect on atmospheric carbon dioxide over the past 100 million years. Am. J. Sci.; (United States) 283:7, Medium: X; Size: Pages: 641–683 2016-2004-2026 (1983). 10.2475/ajs.283.7.641

Boury-Esnault, N., & Rützler. (1997), Thesaurus of sponge morphology. Smithsonian Contributions to Zoology, 596, 65 p. 10.5479/si.00810282.596

Chao, A., Colwell, R. K., Lin, C. W., & Gotelli, N. J. (2009). Sufficient sampling for asymptotic minimum species richness estimators. Ecology, 90, 1125–1133, 10.1890/07-2147.1

Chaudhary, C., Richardson, A.J., Schoeman, D.S., & Costello, M.J. (2021), Global warming is causing a more pronounced dip in marine species richness around the equator. PNAS, 118, e2015094118. 10.1073/pnas.201509411

De Wever, P., O’Dogherty, L., & Gorican, S. (2007), The plankton turnover at the Permo-Triassic boundary, emphasis on radiolarians, in Baumgartner P.O., Aitchison J.C., Wever P., Jackett S-J., eds., Radiolaria: Birkhäuser, Basel, 49–62, 10.1007/978-3-7643-8344-2_4

Jeppsson, L., & Anehus, R. (1995), A buffered formic acid technique for conodont extraction: Journal of Paleontology, 69, 790–794. 10.1017/S0022336000035319

Feng, Q., Yang, F., Zhang, Z., Zhang, N., Gao, Y & Wang, Z. (2000). Radiolarian evolution during the Permian and Triassic transition in South and Southwest China. In: H. Yin., J.M. Dickens., G.R. Shi & J. Tonmg (eds). Persian-Triassic Evolution of Tethys and Western Circum-Pacific. Developments in Palaeontology and Stratigraphy, 18, 309–326.

Foster, W.J., & Twitchett, R.J. (2014), Functional diversity of marine ecosystems after the Late Permian mass extinction event. Nature Geoscience, 7, 233–238. 10.1038/ngeo2079

Foster, W.J., Danise, S., & Twitchett, R.J. (2017), A silicified Early Triassic marine assemblage from Svalbard. Journal of Systematic Palaeontology, 15, 851–877. 10.1080/14772019.2016.1245680

Foster, W.J., Hirtz, J.A., Farrell, C., Reistroffer, M., Twitchett, R.J., Martindale, R.C. (2022). Bioindicators of severe ocean acidifcation are absent from the end-Permian mass extinction. Scientific Reports 12, 1202.

Gates, L., James, N.P., & Beauchamp, B. (2004). A glass ramp: shallow-water Permian spiculitic chert sedimentation, Sverdrup Basin, Arctic Canada. Sedimentary Geology, 168, 125–147.

Godbold, A., Schoepfer, S., Shen, S-Z., & Henderson, C.M. (2017), Precarious ephemeral refugia during the earliest Triassic. Geology, 45, 607–610. 10.1130/G38793.1

Grasby, S.E., & Beauchamp, B. (2009), Latest Permian to Early Triassic basin-to-shelf anoxia in the Sverdrup Basin, Arctic Canada. Chemical Geology 264, 232–246.

Hori, R.S., Yamakita, S., Ikehara, M., Kodama, K., Aita, Y., Sakai, T., Takemura, A., Kamata, Y., Suzuki, N., Takahashi, S., Spörli, B., & Grant-Mackie, J.A. (2011), Early Triassic (Induan) Radiolaria and carbon-isotope ratios of a deep-sea sequence from Waiheke Island, North Island, New Zealand. Palaeoworld, 20, 166–178. 10.1016/j.palwor.2011.02.001

Hounslow, M.W., Peters, C., Mørk, A., Weitschat, W., & Vigran, J.O. (2008), Biomagnetostratigraphy of the Vikinghøgda Formation, Svalbard (Arctic Norway), and the geomagnetic polarity timescale for the Lower Triassic. Geological Society of America Bulletin, 120, p. 1305–1325. 10.1130/B26103.1

Hsieh, T. C., Ma, K. H., & Chao, A. (2016). iNEXT: an R package for rarefaction and extrapolation of species diversity (Hill numbers). Methods in Ecology and Evolution, 7, 1451–1456, 10.1111/2041-210X.12613

Isozaki, Y. (1997), Permo-Triassic boundary superanoxia and stratified superocean: records from lost deep sea. Science, 276, 235–238. 10.1126/science.276.5310.235

Isson, T. T. & Planavsky, N. J. Reverse weathering as a long-term stabilizer of marine pH and planetary climate. Nature 560, 471–475 (2018). 10.1038/s41586-018-0408-4

Isson, T.T., Zhang, S., Lau, K.V., Rauzi, S., Tosca, N.J., Penman, D.E., & Planavsky, N.J. (2022), Marine siliceous ecosystem decline led to sustained anomalous Early Triassic warmth. Nature Communications, 13, 3509. 10.1038/s41467-022-31128-3

Jeppsson, L., & Anehus, R. (1995), A buffered formic acid technique for conodont extraction. Journal of Paleontology, 69, 790–794, 10.1017/S0022336000035319

Kump, L. R. Prolonged Late Permian–Early Triassic hyperthermal: failure of climate regulation? Philosophical Transactions of the Royal Society A: Mathematical, Physical and Engineering Sciences 376, 20170078 (2018)

Lampitt, R.S., Salter, I., and Johns, D. (2009), Radiolaria: Major exporters of organic carbon to the deep ocean: Global Biogeochemical Cycles, 23, GB1010. 10.1029/2008GB003221

Leys, S.P., & Lauzon, N.R. (1998), Hexactinellid sponge ecology: growth rates and seasonality in deep water sponges. Journal of Experimental Marine Biology and Ecology, 230, 111–129. 10.1016/S0022-0981(98)00088-4

Liu, G., Feng, Q., Shen, J.U.N., Yu, J., He, W., & Algeo, T.J. (2013), Decline of siliceous sponges and spicule miniaturization induced by marine productivity collapse and expanding anoxia during the Permian-Triassic crisis in South China. PALAIOS, 28, 664–679. 10.2110/palo.2013.p13-035r

Mørk, A., Elvebakk, G., Forsberg, A.W., Hounslow, M.W., Nakrem, H.A., Vigran, J.O., & Weitschat, W. (1999), The type section of the Vikinghøgda Formation: a new Lower Triassic unit in central and eastern Svalbard. Polar Research, 18, 51–82. 10.3402/polar.v18i1.6558

Muscente, A.D., Prabhu, A., Zhong, H., Eleish, A., Meyer, M.B., Fox, P., Hazen, R.M. & Knoll, A.H. (2018). Quantifying ecological impacts of mass extinctions with network analysis of fossil communities. Proceedings of the National Academy of Sciences, 115, 5217–5222. 10.1073/pnas.1719976115

Nabbefeld, B., Grice, K., Twitchett, R.J., Summons, R.E., Hays, L., Böttcher, M.E., & Asif, M. (2010), An integrated biomarker, isotopic and palaeoenvironmental study through the Late Permian event at Lusitaniadalen, Spitsbergen. EPSL, 291, 84–96. 10.1016/j.epsl.2009.12.053

Nakrem, H. A., & Mørk, A. (1991), New Early Triassic Bryozoa (Trepostomata) from Spitsbergen, with some remarks on the stratigraphy of the investigated horizons. Geological Magazine, 128, 129–140. 10.1017/S001675680001832X

O’Dogherty, L., Carter, E.S., Goričan, Š., & Dumitrica, P. (2010), Triassic radiolarian biostratigraphy. Geological Society, London, Special Publications, 334, 163–200. 10.1144/SP334.8

Penn, J.L., Deutsch, C., Payne, J.L., & Sperling, E.A. (2018), Temperature-dependent hypoxia explains biogeography and severity of end-Permian marine mass extinction: Science, 362, eaat1327. 10.1126/science.aat1327

Racki, G. (1999), Silica-secreting biota and mass extinctions: survival patterns and processes: Palaeogeography, Palaeoclimatology, Palaeoecology, 154, 107–132. 10.1016/S0031-0182(99)00089-9

Reddin, C.J., Aberhan, M., Raja, N.B., & Kocsis, Á.T. (2022), Global warming generates predictable extinctions of warm- and cold-water marine benthic invertebrates via thermal habitat loss. Global Change Biology, 28, 5793–5807. 10.1111/gcb.16333

Ritterbush, K. A., Bottjer, D. J., Corsetti, F. A. & Rosas, S. New evidence on the role of siliceous sponges in ecology and sedimentary facies development in Eastern Panthalassa following the Triassic–Jurassic mass extinction. Palaios 29, 652–668 (2014).

Ritterbush, K. Sponge meadows and glass ramps: state shifts and regime change. Palaeogeography, Palaeoclimatology, Palaeoecology 513, 116–131 (2019)

Sano, H., Kuwahara, K., Yao, A., & Agematsu, S. (2010), Panthalassan seamount-associated Permian-Triassic boundary siliceous rocks, Mino terrane, central Japan. Paleontological Research, 14, 293–314. 10.2517/1342-8144-14.4.293

Sano, H., Kuwahara, K., Yao, A., & Agematsu, S. (2012), Stratigraphy and age of the Permian-Triassic boundary siliceous rocks of the Mino terrane in the Mt. Funabuseyama area, central Japan. Paleontological Research, 16, 124–145. 10.2517/1342-8144-16.2.124

Song, H., Huang, S., Jia, E., Dai, X., Wignall, P.B., & Dunhill, A.M. (2020), Flat latitudinal diversity gradient caused by the Permian–Triassic mass extinction. PNAS, 117, 17578–17583, 10.1073/pnas.1918953117

Takemura, A., Aita, Y., Hori, R.S., Higuchi, Y., Spörli, K.B., Campbell, H.J., Kodama, K., & Sakai, T. (2002), Triassic radiolarians from the ocean-floor sequence of the Waipapa Terrane at Arrow Rocks, Northland, New Zealand. New Zealand Journal of Geology and Geophysics, 45, 289–296. 10.1080/00288306.2002.9514974

Twitchett, R.J., Krystyn, L., Baud, A., Wheeley, J.R., & Richoz, S. (2004), Rapid marine recovery after the end-Permian mass-extinction event in the absence of marine anoxia. Geology, 32, 805–808. 10.1130/G20585.1

Urey, H. C. On the early chemical history of the earth and the origin of life. Proceedings of the National Academy of Sciences 38, 351–363 (1952).

Walker, J. C., Hays, P. & Kasting, J. F. A negative feedback mechanism for the long-term stabilization of Earth’s surface temperature. Journal of Geophysical Research: Oceans 86, 9776–9782 (1981).

Wignall, P.B., Morante, R., & Newton, R. (1998), The Permo-Triassic transition in Spitsbergen: δ13Corg chemostratigraphy, Fe and S geochemistry, facies, fauna and trace fossils. Geological Magazine, 135, 47–62. 10.1017/S0016756897008121

Xiao, Y., Suzuki, N., & He, W. (2017), Water depths of the latest Permian (Changhsingian) radiolarians estimated from correspondence analysis. Earth-Science Reviews, 173, 141–158. 10.1016/j.earscirev.2017.08.012

Yang, F., Sun, Y. D., Frings, P. J., Luo, L., Wang, L. N., Huang, Y. F., Wang T., Müller, J. & Xie, S. C. (2022). Collapse of Late Permian chert factories in the equatorial Tethys and the nature of the Early Triassic chert gap. Earth and Planetary Science Letters, 600, 117861.

Yasuhara, M., Wei, C. L., Kucera, M., Costello, M.J., Tittensor, D.P., Kiessling, W., Bonebrake, T.C., Tabor, C.R., Feng, C.R., Baslega, A., Kretschmer, K., Kusumoto, B., & Kubota, Y. (2020), Past and future decline of tropical pelagic biodiversity. PNAS, 117. 12891–12896. 10.1073/pnas.1916923117

Zuchuat, V., Sleveland, A.R.N., Twitchett, R.J., Svensen, H.H., Turner, H., Augland, L. E., Jones, M.T., Hammer, Ø., Hauksson, B.T., Haflidson, H., Midtkandal, I., & Planke, S. (2020), A new high-resolution stratigraphic and palaeoenvironmental record spanning the End-Permian Mass Extinction and its aftermath in central Spitsbergen, Svalbard. Palaeogeography, Palaeoclimatology, Palaeoecology, 554, 109732. 10.1016/j.palaeo.2020.109732

