## Supplemental Material for "Response of siliceous marine organisms to the Permian-Triassic climate crisis based on new findings from central Spitsbergen, Svalbard"

**Contents of this file**

Text S1

Figures S1

**Paleolatitudes**

| **Region** | **Paleolatitude** | **Reference** |
| --- | --- | --- |
| Sassendalen, Svalbard | 45°N | Hounslow et al. (2008) |
| South China | 8°N | Cai et al. (2014) |
| Mino Terrane, Japan | 6°S | Ando et al. (1986) |
| Arrow Rocks, New Zealand | 36°S | Kodama et al. (2007) |

**LD-01 Thin Section**

B

A

B

A

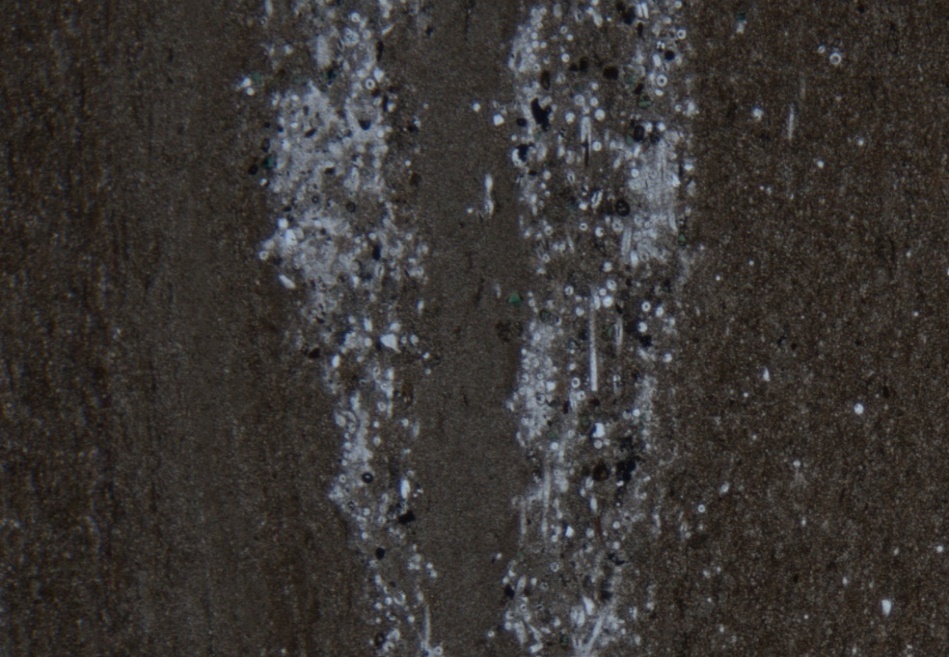

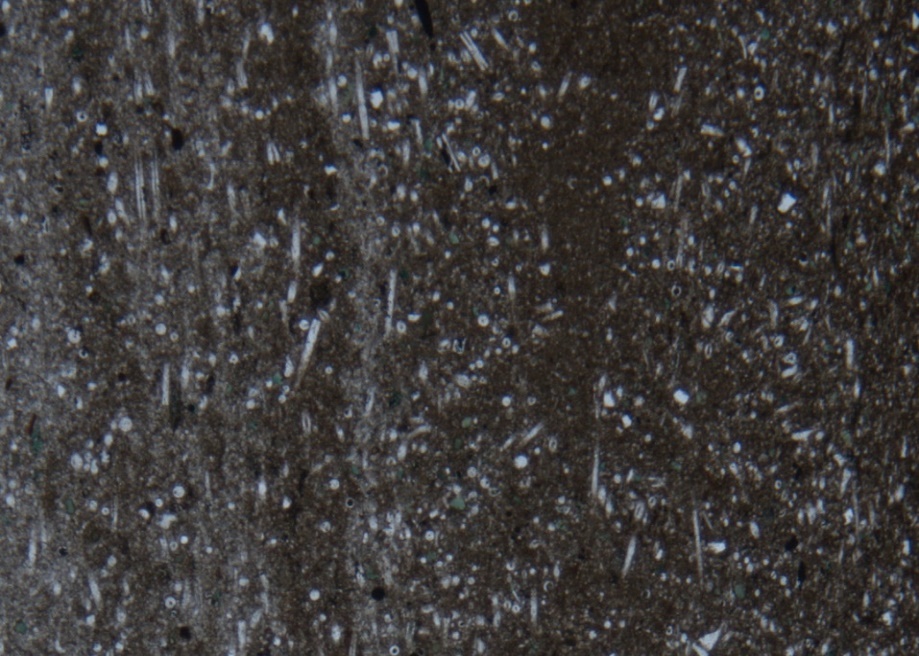

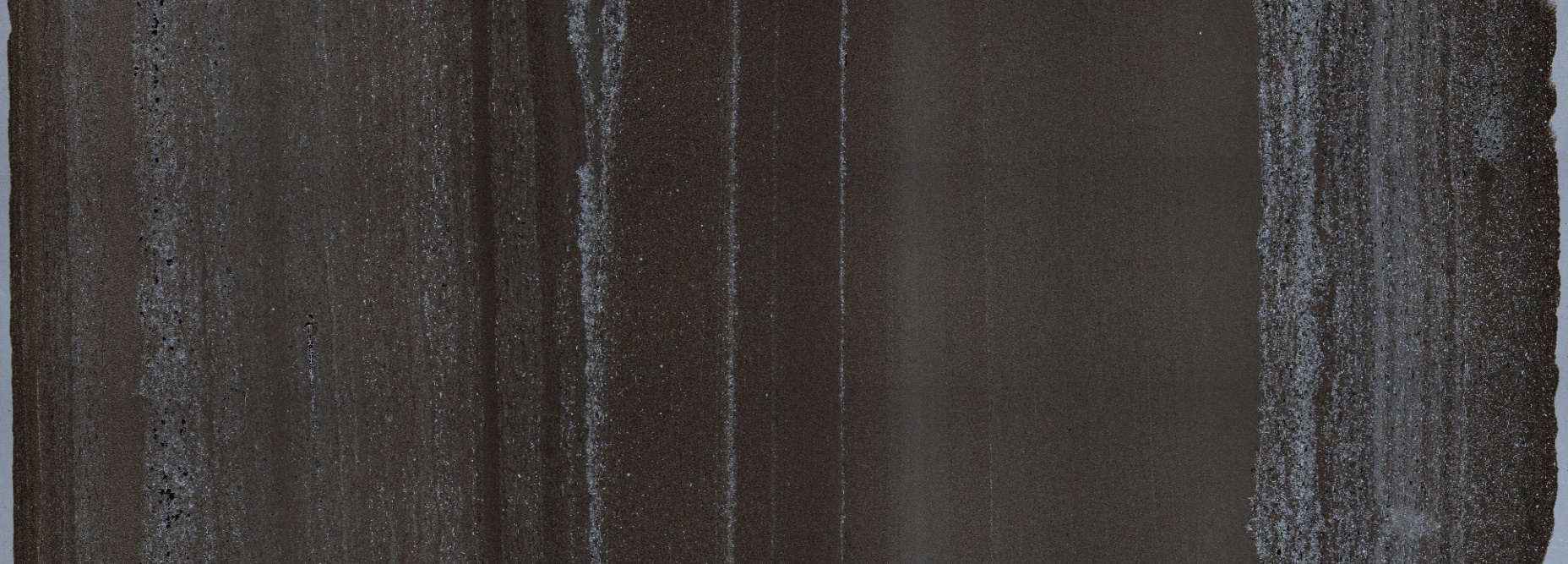

**Figure S1.** Thin section of LD-01 (the first concretionary bed above the extinction horizon) at Lusitaniadalen, Svalbard. A-B) Insets of radiolarian-sponge sub-beds.

**References**

Ando, A., Kodama, K., & Kojima, S. (2001). Low‐latitude and Southern Hemisphere origin of Anisian (Triassic) bedded chert in the Inuyama area, Mino terrane, central Japan. *Journal of Geophysical Research: Solid Earth*, 106, 1973-1986, <https://doi.org/10.1029/2000JB900305>

Cai, J., Tan, X., & Wu, Y., (2014). Magnetic fabric and paleomagnetism of the Middle Triassic siliciclastic rocks from the Nanpanjiang Basin, South China: Implications for sediment provenance and tectonic process. *Journal of Asian Earth Sciences*, 80, 134-147, <https://doi.org/10.1016/j.jseaes.2013.10.033>

Hounslow, M. W., Peters, C., Mørk, A., Weitschat, W., & Vigran, J. O. (2008). Biomagnetostratigraphy of the Vikinghøgda Formation, Svalbard (Arctic Norway), and the geomagnetic polarity timescale for the Lower Triassic. *Geological Society of America Bulletin*, 120, 1305-1325, <https://doi.org/10.1130/B26103.1>

Kodama, K., Fukuoka, M., Aita, Y., Sakai, T., Hori, R. S., Takemura, A., Campbell, H. J., Hollis, C. J., Grant-Mackie, J.A., & Spörli, K. B. (2007). Paleomagnetic results from Arrow Rocks in the framework of paleomagnetism in pre-Neogene rocks from New Zealand, in Spörli, K.B., Takemura, A., and Hori, R.S., eds., The Oceanic Permian/Triassic Boundary Sequence at Arrow Rocks (Oruatemanu), Northland, New Zealand: Lower Hutt, New Zealand. *Geological and Nuclear Science Monograph*, 24, 177-196.

**Table S1.** Radiolarians identified from the basal Triassic (*Hindeodus parvus* Conodont Zone). * = donates Permian holdover species, ^T^ = denotes Triassic range extension, ^P^ = denotes Permian range extension. Data: Svalbard = This Study, Mino terrane, Japan = Sano et al. (2010; 2012), Waiheke Island, New Zealand = Hori et al. (2011), Arrow Rocks, New Zealand = Takemura et al. (2002; 2007).

| **Class** | **Order** | **Family** | **Genus species** | **Svalbard** | **Mino Terrane**  **Japan** | **Waiheke Island**  **New Zealand** | **Arrow Rocks**  **New Zealand** |
| --- | --- | --- | --- | --- | --- | --- | --- |
| Albaillellaria | Albaillellaria | Albaillellidae | *Albaillella* (?) sp. |  |  |  |  |
| Albaillellaria | Albaillellaria | Albaillellidae | *Albaillella* spp. |  |  |  |  |
| Albaillellaria | Albaillellaria | Albaillellidae | *Albaillella aotearoa* |  |  |  |  |
| Albaillellaria | Albaillellaria | Follicucullidae | *Follicucullus* (?) sp. |  |  |  |  |
| Entactinaria | Entactinaria | Entactiniidae | *Copicyntra* sp. |  |  |  |  |
| Entactinaria | Entactinaria | Entactiniidae | *Entactinia itsukaichisensis** |  |  |  |  |
| Entactinaria | Entactinaria | Entactiniidae | *Entactinia* n. sp. |  |  |  |  |
| Entactinaria | Entactinaria | Entactiniidae | *Entactinia nikorni* ^T^ |  |  |  |  |
| Entactinaria | Entactinaria | Entactiniidae | *Entactinia* sp. |  |  |  |  |
| Entactinaria | Entactinaria | Entactiniidae | *Entactinia* cf. *chiakensis* |  |  |  |  |
| Entactinaria | Entactinaria | Entactiniidae | *Entactinosphaera* sp. indet. |  |  |  |  |
| Entactinaria | Entactinaria | Entactiniidae | *Entactinosphaera* sp. |  |  |  |  |
| Entactinaria | Entactinaria | Entactiniidae | *Entactinosphaera spoerlii* |  |  |  |  |
| Entactinaria | Entactinaria | Entactiniidae | *Trilonche crassispinosa* |  |  |  |  |
| Entactinaria | Entactinaria | Entactiniidae | *Oruatemanua* sp. |  |  |  |  |
| Entactinaria | Entactinaria | Pylentonemidae | *Polyentactinia* cf. *phattalungensis* ^T^ |  |  |  |  |
| Entactinaria | Entactinaria | Tetrentactiniidae | *Triaenosphaera minutus* |  |  |  |  |
| Other | Latentifistularia | Cauletellidae | *Cauletella manica* |  |  |  |  |
| Other | Latentifistularia | Cauletellidae | *Cauletella* sp. |  |  |  |  |
| Other | Latentifistularia | Cauletellidae | *Ishigaum* sp. |  |  |  |  |
| Other | Latentifistularia | Cauletellidae | *Pseudotormentus* sp. |  |  |  |  |
| Other | Latentifistularia | Latentifistulidae | *Latentifistula* spp. |  |  |  |  |
| Other | Latentifistularia | Ormistonellidae | *Quadricaulis inflata* |  |  |  |  |
| Other | Latentifistularia | Ormistonellidae | *Quadricaulis scalae* |  |  |  |  |
| Other | Latentifistularia | Pseudolitheliidae | *Grandetortura nipponica* *^P^ |  |  |  |  |
| Other | Latentifistularia | Pseudolitheliidae | *Hegleria* (?) spp. |  |  |  |  |
| Other | Latentifistularia | Pseudolitheliidae | *Hegleria arrowrockensis* |  |  |  |  |
| Other | Latentifistularia | Pseudolitheliidae | *Hegleria mammilla** |  |  |  |  |
| Other | Latentifistularia | Pseudolitheliidae | *Hegleria* sp.* |  |  |  |  |
| Other | Latentifistularia |  | Gen et sp. Indet. (A-B) |  |  |  |  |
| Other | Nassellaria | Spongolophophenidae | *Triassospongocyrtis* (?) sp. |  |  |  |  |
| Other | Nassellaria | Tripedurnulidae | *Tripedocorbis blackae* |  |  |  |  |
| Other | Spumellaria | Actinommidae | *Thaisphaera* (?) *igoi* ^T^ |  |  |  |  |
| Other | Spumellaria | Angulobracchiidae | *Bistarkum martiali* |  |  |  |  |
| Other | Spumellaria | Archaeospongoprunidae | *Archaeospongoprunum* sp. |  |  |  |  |
| Other | Spumellaria | Pyramispongiidae | *Ellipsocopicyntra* sp. |  |  |  |  |
| Other | Spumellaria | Relindellidae | *Copicyntroides* sp. |  |  |  |  |
| Other | Spumellaria | Relindellidae | *Copicyntroides* spp. |  |  |  |  |
| Other | Spumellaria | Sponguridae | *Pseudospongoprunum* (?) spp. |  |  |  |  |
| Other | Spumellaria |  | Gen et sp. indet. (A-F) |  |  |  |  |
